# PURA and GLUT1: Sweet partners for brain health

**DOI:** 10.1101/2023.12.01.569363

**Authors:** Rocío B. Colombo, Clarisa Maxit, Diego Martinelli, Mel Anderson, Diego Masone, Lía Mayorga

## Abstract

PURA, also known as Pur-alpha, is an evolutionarily conserved DNA/RNA-binding protein crucial for various cellular processes, including DNA replication, transcriptional regulation, and translational control. Comprising three PUR domains, it engages with nucleic acids and has a role in protein-protein interactions. The manifestation of PURA syndrome, arising from mutations in the PURA gene, presents neurologically with developmental delay, hypotonia, and seizures. In our prior work from 2018, we highlighted the unique case of a PURA patient displaying hypoglycorrhachia, suggesting a potential association with GLUT1 dysfunction in this syndrome.

In this current study, we expand the patient cohort with PURA mutations exhibiting hypoglycorrhachia and aim to unravel the molecular basis of this phenomenon. We established an in vitro model in HeLa cells to modulate PURA expression and investigated GLUT1 function and expression. Our findings indicate that PURA levels directly impact glucose uptake through the functioning of GLUT1, without influencing significantly GLUT1 expression. Moreover, our study reveals convincing evidence for a physical interaction between PURA and GLUT1, demonstrated by colocalization and co-immunoprecipitation of both proteins. Computational analyses, employing molecular dynamics, further corroborates these findings, demonstrating that PURA:GLUT1 interactions are plausible, and that the stability of the complex is altered when PURA is truncated and/or mutated.

In conclusion, our results suggest that PURA plays a pivotal role in driving the function of GLUT1 for glucose uptake, potentially forming a regulatory complex. Additional investigations are warranted to elucidate the precise mechanisms governing this complex and its significance in ensuring proper GLUT1 function.

## INTRODUCTION

The protein PURA, also known as Pur-alpha, is a highly conserved single-stranded DNA/RNA binding protein with intricate involvement in various cellular processes, including DNA replication, RNA processing, transcriptional and translational regulation[1]. Since its initial characterization in 1992 in HeLa cells [2], extensive research has revealed its multifaceted roles in modulating the functions of numerous molecules across different cellular compartments.

As a member of the PUR family, PURA comprises three highly conserved sequence-specific repeats known as PUR domains I, II, and III, which play a fundamental role in the protein’s primary functions[3]. Each PUR amino acid repeat is characterized by a structural configuration featuring a β-sheet domain and an α-helical domain organized in a “whirly fold” arrangement. In this structure, the convex β-sheets provide a surface for interactions with nucleic acids, while the remaining helical portions are actively engaged in protein-protein interactions[4]. Regarding protein-protein interactions, specifically, the PUR I and II motifs are suggested to establish intramolecular peptide-peptide bonds with each other. In contrast, the PUR III domain assumes the responsibility of facilitating homo-heterodimerization with another PUR protein or interacting with other proteins[3,5].

Although initially described as primarily localized in the nucleus, PURA has demonstrated pivotal functions spanning various cellular domains. Within the nucleus, PURA binds to distinct DNA promoter regions of genes, such as MBP[6], CD43[7–10], and GATA2[11]. In addition, it can collaborate synergistically with transcription factors like E2F1[12,13]. Furthermore, it exhibits binding affinity towards circular RNAs [14] and non-coding RNAs[15], exerting notable influences on gene transcription. Additionally, PURA plays a significant role in DNA repair processes[16,17].

In the cytoplasm, PURA’s capabilities extend to mRNA binding, promoting site-specific translation and participating in the formation of RNA binding protein (RBP) granules, including but not limited to BCL1, MAP2, Stau1, Myo5A, and FMR1 [18,19]. Direct protein-protein interactions have been observed with KIF1[20] and other members of the PUR family[4,21].

Notably, PURA emerges as a crucial player in disease scenarios, particularly in conditions involving CGG repeat-expanded FMR1 (Fragile-X syndrome) [22,23] and G_4_C_2_ repeat expansions in C9orf72, contributing to Amyotrophic Lateral Sclerosis and Frontotemporal degeneration (ALS/FTD) [24,25]. Here, it binds to anomalous RNAs, serving a protective role against neurotoxicity.

The advent of next-generation sequencing and high-throughput genetic technologies in clinical practice has led to increased identification of patients with PURA syndrome[26–35]. PURA syndrome, a rare genetic condition resulting from mutations in the PURA gene, manifests as a spectrum of clinical features, mainly neurological, including developmental delay, hypotonia, breathing difficulties, pituitary dysfunction, and seizures[26–35]. Evidently, PURA’s indispensable role in neuronal development and function is underscored by these clinical findings in patients and by evidence gained from two knock-out mouse models[36,37], which exhibited severe tremors, seizures, movement disorders associated with alterations in brain size, cell distribution and premature death. Intriguingly, one of the knock-out models was pointed out to be able to extend their short lifespan with periodical glucose injections[36]. Furthermore, human patients with PURA syndrome, have been described to manifest with hypoglycemia[30] introducing a possible link between PURA and glucose metabolism.

The transportation of glucose across the blood-brain barrier predominantly relies on the glucose transporter 1 (GLUT1 or SLC2A1), highly expressed in endothelial cells, astrocytes and red blood cells[[38]. Mutations in GLUT1 give rise to GLUT1 deficiency syndrome, characterized by seizures, developmental delay, and movement disorders, accompanied by the disordeŕs hallmark: hypoglycorrhachia (low cerebrospinal fluid glucose levels) [39,40]. Notably, PURA syndrome and GLUT1 deficiency share numerous neurological clinical features, but hypoglycorrhachia was first reported by us in 2018 in a PURA syndrome patient[35]. This discovery suggested a potential connection between PURA and GLUT1. In this current study, we expand the cohort of patients suffering from PURA deficiency, who also displayed hypoglycorrhachia. This intriguing observation served as the cornerstone of our research, laying the foundation for our findings that establish a direct link between PURA and GLUT1.

## METHODS

### Patient data

PURA deficiency patientś data were gathered from medical records from the care-giving centers and physicians.

### Cell culture

Human HeLa (CCL-2) cells were purchased from ATCC, The Global Bioresource Center and grown in DMEM high-glucose (Gibco®, cat#11995065) in 5% CO2 supplemented with 10% fetal bovine serum (Gibco®, cat#16000044), 100U/ml Penicilin+ Streptomicine (Gibco®, cat#15140122). Passages 20-38 were used for experiments. The six experimental conditions used for experiments were created as follows:

1_ Control. Generation of a stable cell line of HeLa with the mammalian IPTG-Inducible shRNA PURA knockdown lentiviral vector, purchased in Vector Builder® pLV[shRNA]-LacI:T2A:Puro-U6/2xLacO>hPURA[shRNA#1]. Lentivirus were produced in Hek293T cells by transient co-transfection of the MISSION® Lentiviral Packaging Mix (Sigma®) with 1.35µg of the shRNA-encoding DNA using 7 µg polyethylenimine (PEI, Poliscience®) for transfection of a well from a 6-well plate. Culture supernatants were collected 72 h post-transfection, filtered (0.22 mm filter, cellulose acetate, Ministart®), and supplemented with 8 μg/ml polybrene (Sigma®). Subconfluent HeLa cells ̴50.000 in a well from a 6-well plate were infected with the polybrene-supplemented supernatant and 48h post-infection were selected with 1 μg/ml puromycin (sc-108071®) for 1 week.

2_shPURA (IPTG-induced knock-down). HeLa cells from condition 1 were treated with IPTG: isopropyl-galactosidase (TRANS®) 500μM for 72h (replacing medium and IPTG every day)

3_+PURA: PURA overexpression: ∼50.000 HeLa cells from condition 1 were transfected with 1.5µg VB220517-1200nex pRP[Exp]-mCherry/Neo-CMV>hPURA[NM_005859.5] purchased in Vector Builder® using 7 µg polyethylenimine (PEI, Poliscience®). 12hours after, medium was replaced and cells were selected through Geneticin sulfate (G 418_ab144261) 200 µg/ml for 72h.

4_ +GLUT1 (GLUT1 overexpression): ̴50.000 HeLa cells from condition 1 were transfected with 1.5 µg pRP[Exp]-Bsd-CAG>hSLC2A1[NM_006516.4] purchased in Vector Builder® using 7 µg polyethylenimine (PEI, Poliscience®). 12hours after, medium was replaced and cells were selected through Blasticidin S hydrochloride, Nucleoside (ab141452) 10 µg /ml for 72h.

5_+GLUT1+shPURA (concomitant GLUT1 overexpression and induced PURA knockdown): cells were treated as condition 4 and during the selection process IPTG was added as in condition 2.

6_+PURA+GLUT1 (PURA and GLUT1 co-overexpression). A combination of condition 3 and 4 was carried out. Both plasmids were transfected simultaneously, and selection was carried out with both antibiotics (Geneticin+ Blasticidin) for 72h.

### Glucose uptake assays

HeLa cells from the 6 conditions above, on their second day of selection were seeded in glass coverslips. 24h later, 2-(N-(7-nitro-2,1,3-benzoxadiazol-4-yl)amino)-2-deoxyglucopyranoside (2-NBDG, Medkoo®) was added in DMEM low glucose medium to a 30µM concentration for 1h. Then, cells were washed with PBS and imaged in DMEM FluoroBrite medium using Zeiss Axio Observer fluorescence inverted microscope® or tripsinized and analyzed through BD Accuri C6 Plus Flow Cytometer®, fluorescence being evaluated with channel FL1 (533/30nm). Microscopy fluorescence was evaluated using Image J® software (mean grey value from +50 cells per image ̴5 photos per condition substracting background). Flow cytometry assays were analyzed using FloJo v X.0.7® software.

### mRNA expression, RTqPCR

RNA was extracted from HeLa cells (conditions 1,2,3) using TriPure® (Roche®) according to manufacturerś protocol. RNA retrotranscription to obtain cDNA was performed using M-MLV retrotranscriptase (Inbio Highway®) based on manufactureŕs protocol. SYBR based qPCR for genes: PURA, GLUT1 and ACTINB or B2M as housekeeping genes was performed with SsoAdv univer SYBR GRN BioRad®. PURA and GLUT1 reaction parted from 40ng of RNA derived cDNA, housekeeping geneś reaction from 10ng. Primer concentration for all 4 pairs was 250nM of each. qPCR was carried out with the following thermal program: 95 °C 3 min; 40 cycles of: 95 °C 15 s, 60 °C 30 s (read), 72 °C 30 s using the BioRad PCR CFX96® qPCR equipment. Results were analyzed with CFX Maestro Software®. Quantification was estimated through ΔΔCT method.

Primerś sequences:

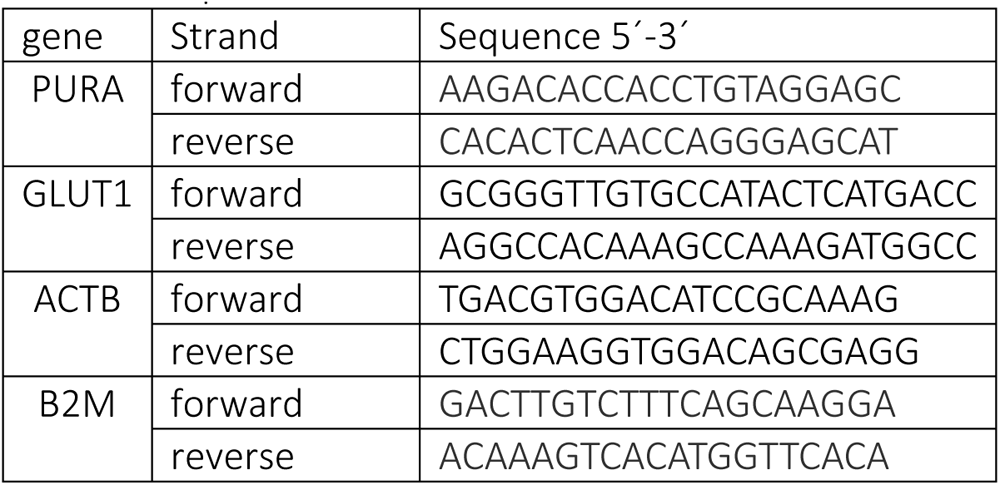

### Western Blots

Cells were lysed with lysis buffer solution compatible with Co-Immunoprecipitation assays: 1mM EDTA, 10% glycerol, 75mM NaCl, 0.05% SDS, 100mM Tris-Cl pH 7.4, 0.1% (v/v) triton X-100) and measured through BCA kit (Thermofisher Scientific®). Proteins (10–30 μg) were run on a 10% SDS–polyacrylamide gels and then transferred to a nitrocellulose membrane (Biorad®). The membranes were blocked in 5% lowfat milk-PBS-tween solution and then incubated overnight at 4 °C with primary antibodies: anti-PURA 1:1500 (rabbit—ab125200 abcam®), anti-GLUT1 1:2000 (mouse— [GLUT1/2476] (A249979) Antibodies®) and anti-Vinculin Antibody 1:2000 CAB2752-20 (rabbit—GENIE®) followed by PBS-Tween washes and secondary antibody incubation (2h at room temperature): Horseradish peroxidase secondary antibodies: mouse (Gt anti-mouse IgG -H+L-Invitrogen®) 1:10000 and rabbit (Gt anti-Rb IgG -H+L-Invitrogen®) 1:3000. Bands were developed using chemiluminescence (ECL BPS-Bioscience®), visualized with a LAS Fujifilm 4000 system (GE Healthcare Life Sciences®) and quantified using Image J® software.

### Indirect immunofluorescence

Cells were fixed with 4% Paraformaldehyde solution 30 min at room temperature, following washes with PBS, and then quenched with ClNH_4_ 50 mM for 30 min at room temperature. Cells were permeabilized with Albumin 2%/Saponin0.1% PBS solution, following which they were incubated with anti-PURA antibody (rabbit—ab125200 abcam®) 1:200 and anti-GLUT1 (mouse-RyD MB14181®) 1:100 or [GLUT1/2476] (A249979) Antibodies®) 1:200, overnight at 4 °C. The primary antibody was rinsed and then cells were incubated with secondary antibodies ab150077 Goat Anti-Rabbit IgG H&L (Alexa Fluor® 488) 1:750 and ab150115 Goat Anti-Mouse IgG H&L (Alexa Fluor® 647) 1:750, washed, and coverslips were mounted on glass slides using Mowiol + Hoescht (LifeTechnologies®) and examined by fluorescence confocal microscopy (Olympus Confocal Microscope FV1000-EVA®).

### Dot blot overlay

A nitrocellulose membrane was imbedded with 1µl of antibodies targeting PURA (rabbit-ab125200 abcam®) or GLUT1 (mouse-RyD MB14181®), blocked with BSA 5% solution 1h room temp, then the exposed to ̴ 40µg HeLa cell lysates overnight at 4 °C. Membranes were washed and subsequently exposed to the counterpart proteińs antibody overnight at 4 °C (nitrocellulose membrane that was imbedded with anti-PURA was incubated with anti-GLUT1 and vice versa, 1:500 for anti-PURA and 1:250 for anti-GLUT1 antibodies). Membranes were washed and then incubated with Horseradish peroxidase secondary antibodies against the last antibody: mouse (Gt anti-mouse IgG -H+L-Invitrogen®) 1:10000 and rabbit (Gt anti-Rb IgG -H+L-Invitrogen ®) 1:3000. Dot blots were developed using chemiluminescence (ECL BPS-Bioscience®) and visualized with a LAS Fujifilm 4000 system (GE Healthcare Life Sciences®).

### Co-Immunoprecipitation

Immunoprecipitation was carried out through ImmunoPure Plus Immoblized Protein A agarose beads (PIERCE®) bond to anti-PURA ab125200 abcam® following protocol available at https://www.abcam.com/protocols/immunoprecipitation-protocol-1. We followed method B from the mentioned protocol using 15µl of bead slurry + 0.5 µg of anti-PURA antibody per 30 µg of protein from cell lysate. After thorough washes, beads harboring the pulldown were run on a 10% SDS–polyacrylamide gel and the presence of GLUT1 was analyzed through Western Blot. RNAse A (Thermofisher Scientific®) was used to evaluate RNA dependance of the protein complex PURA:GLUT1. We used 0.5µg RNAse A per 12µl of lysate, 10min 25°C incubation prior to the immunoprecipitation protocol.

### Computational methods

#### Atomistic molecular dynamics

Atomistic simulations of GLUT1 and PURA in complex were conducted with Gromacs-2023[41]. Proteins were solvated using CHARMM-GUI web server[42]. All protein-protein complexes were minimized and equilibrated following the CHARMM-GUI protocol for Gromacs[43]. For productions runs the pressure and the temperature were set to 1.0 bar and 303.15K. Pressure was controlled by the Parrinello-Rahman barostat[44].Temperature was controlled by de Nose-Hoover thermostat[45] using a 1ps coupling constant. Cut-offs for van der Waals and electrostatic interactions were set to 1.2nm, using the particle Mesh Ewald (PME) mehtod[46] with Verlet cut-off scheme. In all cases a 4^th^ order LINCS algorithm was applied[47].

### Article drafting

we used ChatGPT, an AI-powered language model developed by OpenAI, solely for language-related assistance in the composition of this research paper, with no influence on the content or research outcomes.

### Statistics software

Graph Pad Prism v5.03® was used for statistical analyses and graph confection.

### Art work

Corel draw 2020® was used to draft figures. Computational figures were prepared with Visual Molecular Dynamics (VMD)[48], Schrödinger academic version, MDVWhole, Grace (GRaphing, Advanced Computation and Exploration of data) and Inkscape. Biorender® was used to graph the plasma membrane in figure 3.

## RESULTS

### Hypoglycorrhachia: a recurrent finding in PURA syndrome patients

In 2018, our research team reported the inaugural case of a patient exhibiting PURA deficiency with the distinctive manifestation of hypoglycorrhachia[35]. Notably, this patient displayed diminished GLUT1 expression within blood cells. In the present study, we expand upon our previous work by presenting data from an additional cohort of patients, who exhibited as well, the hallmark symptom of hypoglycorrhachia. A comprehensive summary of the key clinical observations for these patients, along with details regarding PURA mutations and the corresponding glycorrhachia levels, as well as the glycorrhachia-to-glycemia ratio, is presented in Table 1.

**Table 1.** presents a cohort of individuals diagnosed with PURA syndrome manifesting hypoglycorrhachia. The table includes information on DNA and mRNA mutations, protein consequences, glycorrhachia levels, and the corresponding glycorrhachia/glycemia ratios. Additionally, the table summarizes key clinical features associated with the presented cases.

| patient | PURA mutation (heterozygosis and *de novo*) Human (GRCh38/hg38) genome | PURA mutant protein | Mean Glycorrachia (normal value >2.9mM)[40] | Mean Glycorrachia-to-glycemia ratio (normal value>0.59)[40] | Other clinical aspects | Care center location | Previously reported |
| --- | --- | --- | --- | --- | --- | --- | --- |
| 1 | Chr5: 140114767delA<br>NM_005859.4 c.586delA | p.I196fs*30 (223 aminoacids) | 1.6mM | 0.39 | Hypotonia, seizures, apnea, development delay | Mendoza, Argentina | Yes [30,35] |
| 2 | Chr5:<br>140114918del<br>TGAGCGA<br>NM_005859.4<br>c.737_744del | p.V246<br>Gfs*45<br>(289 aa) | 2.9mM | 0.44 | Hypotonia,<br>seizures,<br>development delay | Buenos<br>Aires<br>,<br>Argentina | No |
| 3 | Chr5:<br>140114992del<br>TTC<br>NM_005859.4<br>c.810_812<br>delTTC | p.F271<br>del<br>(321aa) | 2.1mM | 0.55 | Hypotonia,<br>seizures,<br>apnea,<br>development delay | Rome,<br>Italy | No |
| 4 | Chr5:<br>140114714dup<br>TGGCCTGGGCT<br>CC<br>NM_005859.4<br>c.534_546dup<br>TGGCCTGGGCT<br>CC | p.T183<br>Wfs*22<br>(203aa) | 2.2mM | 0.44 | Hypotonia,<br>seizures,<br>development delay | Rome,<br>Italy | No |
| 5 | Chr5:<br>140114386C>T<br>NM_005859.4<br>c.205C>T | p.Q69*<br>(68aa) | 2.3mM | 0.42 | Hypotonia,<br>seizures, development delay | Rome,<br>Italy | No |
| 6 | Chr5:<br>140114778del<br>TTC<br>NM_005859.4<br>c.697_699<br>delTTC | p.F233<br>del<br>(321aa) | 2.2mM | 0.47 | Hypotonia,<br>seizures,<br>development delay,<br>hyperprolactinemia | Melbourne,<br>Australia | Yes[26,30] |

An intriguing observation stemming from our analysis is the consistent association between hypoglycorrhachia and specific PURA mutations affecting the PUR III repeat region within the protein’s structure domain which is believed to facilitate interactions with other proteins [3]. In light that hypoglycorrhachia could now be considered a new finding in patients with PURA deficiency and observing that the rest of the clinical features overlap, we ventured to take these clinical findings to the bench side and investigate a functional connection between PURA and GLUT1.

### PURA enhances glucose uptake in cultured HeLa cells by potentiating the function of GLUT1, without inducing significant alterations in GLUT1 expression levels

Firstly, we wanted to prove in an *in vitro* model a functional interplay between these proteins. Accordingly, glucose uptake served as the ultimate metric to assess the influence of PURA on GLUT1’s functionality.

PURA was originally delineated in HeLa cells[2], and as a result, we devised our experimental setup utilizing these cells as our chosen model system. We established a stable HeLa cell line containing an inducible shRNA targeting the PURA gene. These cells were manipulated to either suppress PURA expression (via IPTG shRNA induction) or augment PURA expression (via transient transfection with a PURA expression plasmid). In addition, we used GLUT1 overexpression as a positive control. To discern the critical role of PURA in glucose uptake and its interrelationship with GLUT1, we devised six distinct conditions as follows: 1_Control (PURA baseline), 2_ shPURA (IPTG-induced PURA knockdown), 3_ +PURA (PURA overexpression), 4_ +GLUT1 (GLUT1 overexpression), 5_+GLUT1+shPURA (concomitant GLUT1 overexpression and induced PURA knockdown), 6_+PURA+GLUT1 (PURA and GLUT1 co-overexpression). We validated the precision of the proposed manipulations by Western Blot. (Fig.1A)

To evaluate glucose uptake across these six conditions, we employed a fluorescent glucose analog, 2-deoxy-2-[(7-nitro-2,1,3-benzoxadiazol-4-yl)amino]-D-glucose (2-DNBG). Subsequently, we quantified fluorescence levels utilizing both fluorescent microscopy (Fig.1B) and flow cytometry techniques (Fig. 1C).

Our findings revealed that PURA upregulation significantly enhanced glucose uptake, comparable to the impact of sole GLUT1 overexpression, as visually depicted in microscopy (Fig.1B, condition 3) and vindicated through flow cytometry (Fig.1C). Conversely, PURA knockdown exerted a diminishing effect on glucose uptake, evident in basal conditions as well as in cells with heightened GLUT1 expression, as illustrated in microscopy (Fig.1B, conditions 2 and 5). These experiments show that PURA is a major exponent for glucose uptake and particularly condition 5’s result (diminished glucose uptake when PURA is knocked down in a high GLUT1 cellular environment) shows a functional interdependence of GLUT1 towards PURA, indicating a solid interplay between these two proteins. In addition, the absence of synergy in glucose uptake under condition 6 (overexpression of both proteins) indicates that PURA and GLUT1 function in the same pathway for glucose incorporation.

Given PURA’s established capability to modulate gene transcription and translation, and the finding of diminished GLUT1 expression in blood cells from a patient with PURA syndrome[35], firstly, we thought that PURA might regulate GLUT1’s transcription and/or translation. To evaluate this, we conducted a focused examination of GLUT1 mRNA and protein expression levels during instances of PURA up and downregulation (Fig. 1D). Notably, these experimental manipulations did not yield significant alterations in GLUT1 levels, leaving us to investigate other possibilities of partnership between the proteins. Taking into account that the PUR III repeat domain is known to facilitate interactions with other proteins[3], what if PURA and GLUT1 were to interact in a complex?

**Figure 1:**
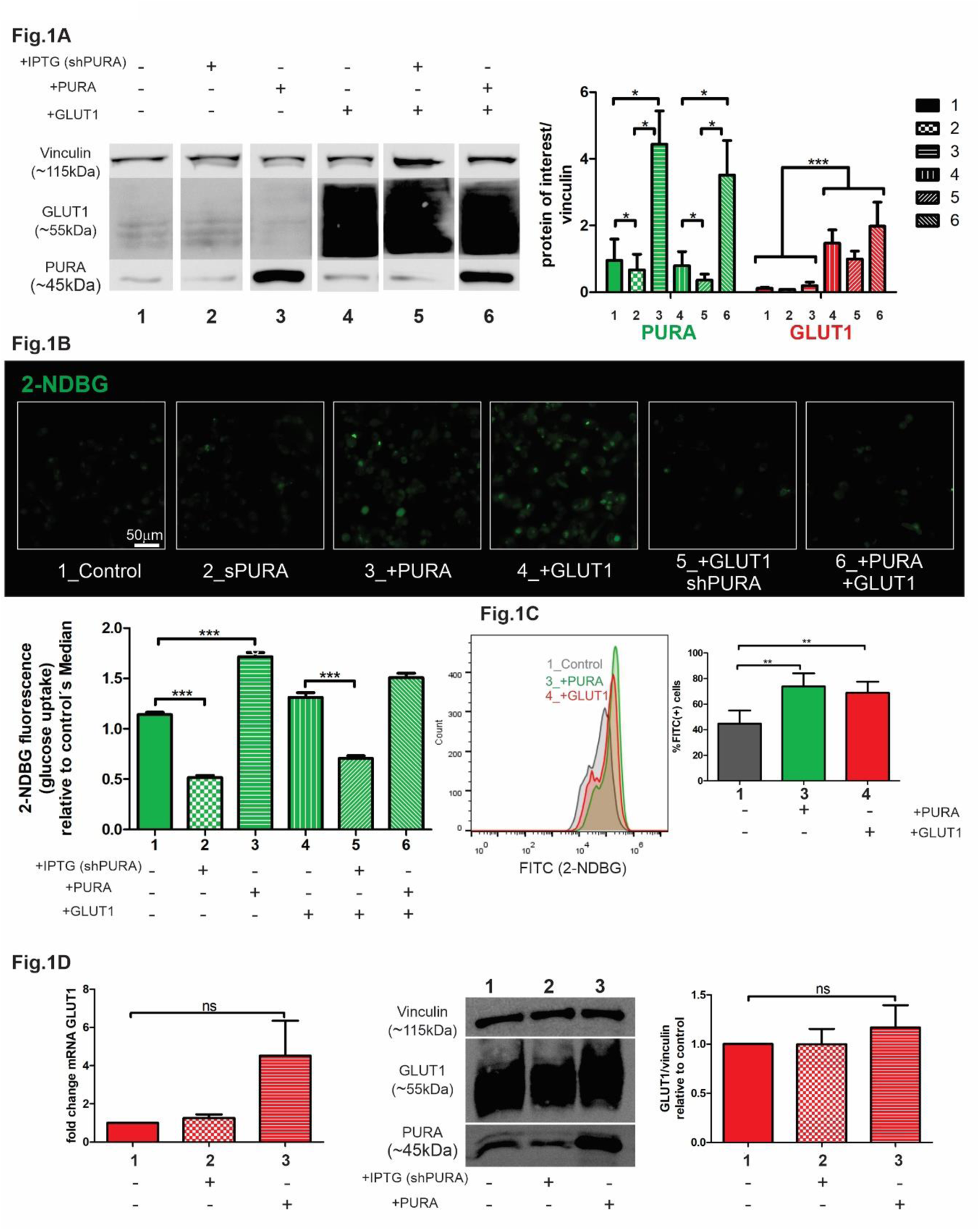
PURA enhances glucose uptake in cultured HeLa cells without inducing significant alterations in GLUT1 expression levels. **1A**. Validation by Western Blot of the proposed expression manipulation of PURA and GLUT1. 1_Control (PURA baseline), 2_ shPURA (IPTG-induced PURA knockdown), 3_ +PURA (PURA overexpression), 4_+GLUT1 (GLUT1 overexpression), 5_+GLUT1+shPURA (concomitant GLUT1 overexpression and induced PURA knockdown), 6_+PURA+GLUT1 (PURA and GLUT1 co-overexpression). All lanes correspond to the same gel but have been cut in order to follow the 1 to 6 conditionś order N=3. PURA*p<0.05. One-tailed paired student’s T-test; GLUT1: ***p<0.001 Two-tailed paired Student’s T test between all conditions with basal GLUT1 level (conditions 1,2,3) and all conditions with GLUT1 overexpression (conditions 4,5,6). **1B and C**. Glucose uptake assay using the fluorescent glucose analog, 2-deoxy-2-[(7-nitro-2,1,3-benzoxadiazol-4-yl)amino]-D-glucose (2-NDBG). We quantified fluorescence levels with fluorescent microscopy (**1B**: Glucose uptake was different in all six conditions except 3 vs 6, Kruskal-Wallis test, Dunńs Multiple Comparison post-test. Significance was p<0.01 for all significant comparisons. We have selectively displayed on the graph only those comparisons that are most pertinent to convey the core message of this paper, ***p<0.001); and flow-cytometry (1C, **p<0.01 One-way ANOVA+ Tukey’s Multiple Comparison Test). **1D**. GLUT1 expression was measured at an mRNA level through SYBR based qPCR, Δ ΔCT quantification method (left panel) and at a protein level through Western Blot (right side), One-way ANOVA+ Tukey’s Multiple Comparison Test. ns: not significant.

### PURA and GLUT1 are observed to engage in a cellular complex independently of RNA interactions

In light of the fact that the functional association between PURA and GLUT1 did not appear to hinge on the abundance of GLUT1, we embarked on investigating the potential formation of a physical complex between these two proteins.

Through our experiments we revealed several lines of evidence that support the notion of a physical interaction between PURA and GLUT1:

Immunofluorescence microscopy revealed co-localization of PURA and GLUT1 within HeLa cells (Fig. 2A). This co-localization was more pronounced when both proteins were concurrently overexpressed or when PURA overexpression was coupled with indirect GLUT1 upregulation with rotenone (mitochondrial complex 1 inhibitor) treatment 10µM 24h(mean Pearsońs coefficient 0.81: Supplementary material Fig. S1A and S1B). [49,50].

Dot-blot overlay assays yielded additional evidence of the interaction between PURA and GLUT1. A nitrocellulose membrane was imbedded with the antibody targeting PURA (Fig 2B., left panel) or GLUT1 (Fig. 2B, right panel) and then exposed to HeLa cell lysates overexpressing PURA. As a result, these antibodies effectively captured and immobilized the respective counterpart protein upon the nitrocellulose membrane, adding further evidence for PURA-GLUT1 interaction (Fig. 2B).

Co-immunoprecipitation experiments were conducted using anti-PURA antibody conjugated to Protein A agarose beads. Proteins from cell lysates overexpressing PURA were exposed to the immune-anti-PURA-coated beads. The pull-down was isolated and subsequently analyzed by Western Blot, confirming the presence of GLUT1 in the pull-down (Fig. 2C, left panel). Importantly, this interaction persisted even when PURA levels were reduced through knockdown, providing compelling evidence for its persistence under conditions of diminished PURA expression (Supplementary Material Fig. 2S). Accordingly, we can affirm that PURA and GLUT1 interact with each other. However, this interaction could be direct or indirect.

Considering PURA’s well-established role as an RNA-binding protein primarily involved in the formation of ribonucleoprotein (RNP) complexes, we asked ourselves if this interaction could be mediated by RNA. Therefore, we introduced a condition wherein lysates were pretreated with RNAse A. Remarkably, this treatment did not alter the quantity of GLUT1 detected in the pulled-down complexes (Fig. 2B, right panel). This finding suggests an RNA-independent interaction between PURA and GLUT1.

**Figure 2:**
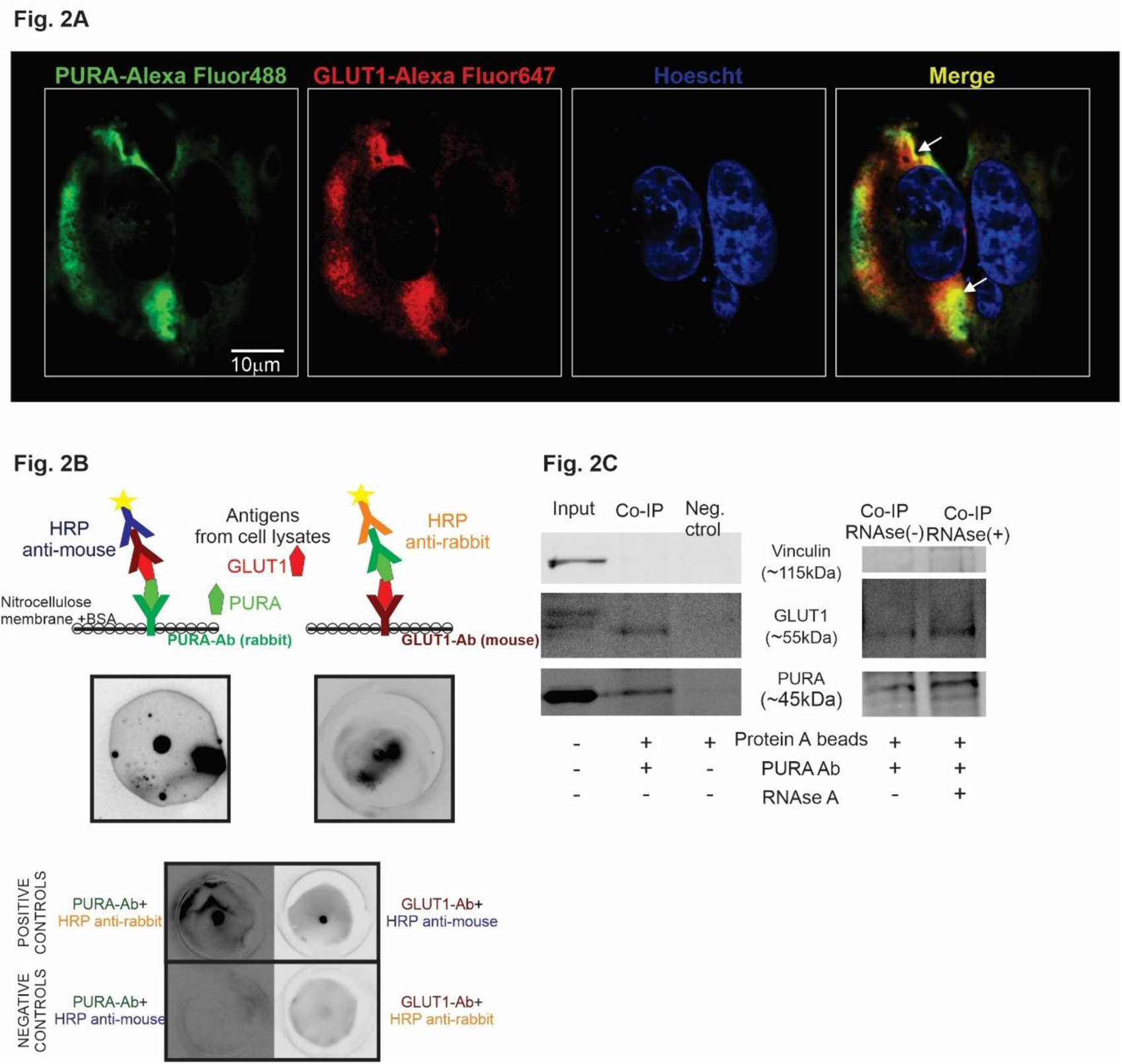
PURA and GLUT1 form a complex. **2A**: Confocal Immunofluorescence Microscopy. Confocal laser micrographs depict indirect immunofluorescence staining of PURA (labeled with AlexaFluor 488 in green) and GLUT1 (labeled with AlexaFluor 647 in red) within HeLa cells overexpressing PURA and treated with Rotenone to enhance GLUT1 expression. The cell nuclei were stained with Hoescht (blue). Colocalization of the two proteins in the merged image is represented as yellow, indicated by white arrows. **2B**: Dot Blot Overlay Assay. PURA (left panel) and GLUT1 (right panel) antibodies were imbedded in the center of circular pieces of nitrocellulose membrane, BSA was used to block non-specific interaction. HeLa cell lysates from cells overexpressing PURA were exposed to the membrane and subsequently washed. Following this, the membrane was incubated with antibodies targeting the counterpart protein, followed by secondary antibody incubation with horseradish peroxidase (HRP) reaction visualization. Controls are shown in the bottom of the figure. Positive and negative controls were processed similarly, excluding incubation with cell lysates, to demonstrate antibody specificity. Notably, PURA antibodies are of rabbit origin, while GLUT1 antibodies are of mouse origin, allowing for their differentiation based on the secondary antibody’s signal. **2C**. Co-immunoprecipitation of GLUT1 and PURA using Protein A agarose beads bond to anti-PURA antibody. The left panel demonstrates the presence of GLUT1 in the pull-down fraction associated with Protein A agarose beads and anti-PURA antibody (Ab), while it is absent in the pull-down from beads lacking anti-PURA Ab. The right panel reveals that treatment with RNAse does not influence the pull-down of GLUT1 facilitated by PURA-Protein A beads, suggesting that their interaction is not mediated by RNA.

### PURA mutations identified in patients displaying hypoglycorrhachia were found to exhibit altered protein-protein affinity, as through computational analysis

With no crystallographic structure still available for PURA (Uniprot: Q56A79), we used AlphaFold[51] as a starting point for its computational structural prediction. Figure 3 shows the resulting PURÁs structure (res 57-322), the first 56 residues are predicted to be disordered and were deleted from the figure to make it visually more comprehensive. Additionally, to account for the PDZ binding motif in GLUT1 (res 488-492, DSQV), we also used AlphaFold to predict missing residues in the original crystal structure of the human sugar transporter GLUT1 (PDB ID: 6THA) [52,53]. Figure 3 shows full-length GLUT1 (res 1-492).

PURÁs unstructured N-terminal was deleted before computational analysis. Initial binding sites for docking in PURA and GLUT1 were selected at the middle of Repeat-III region (res 270-300) and at the C-terminal PDZ binding motif (res 488-492), respectively and according to previous experimental studies that predict the PURA PURIII domain and the GLUT1 PDZ domain as the main protein-protein interacting regions[3,54] (Fig 3).

**Figure 3:**
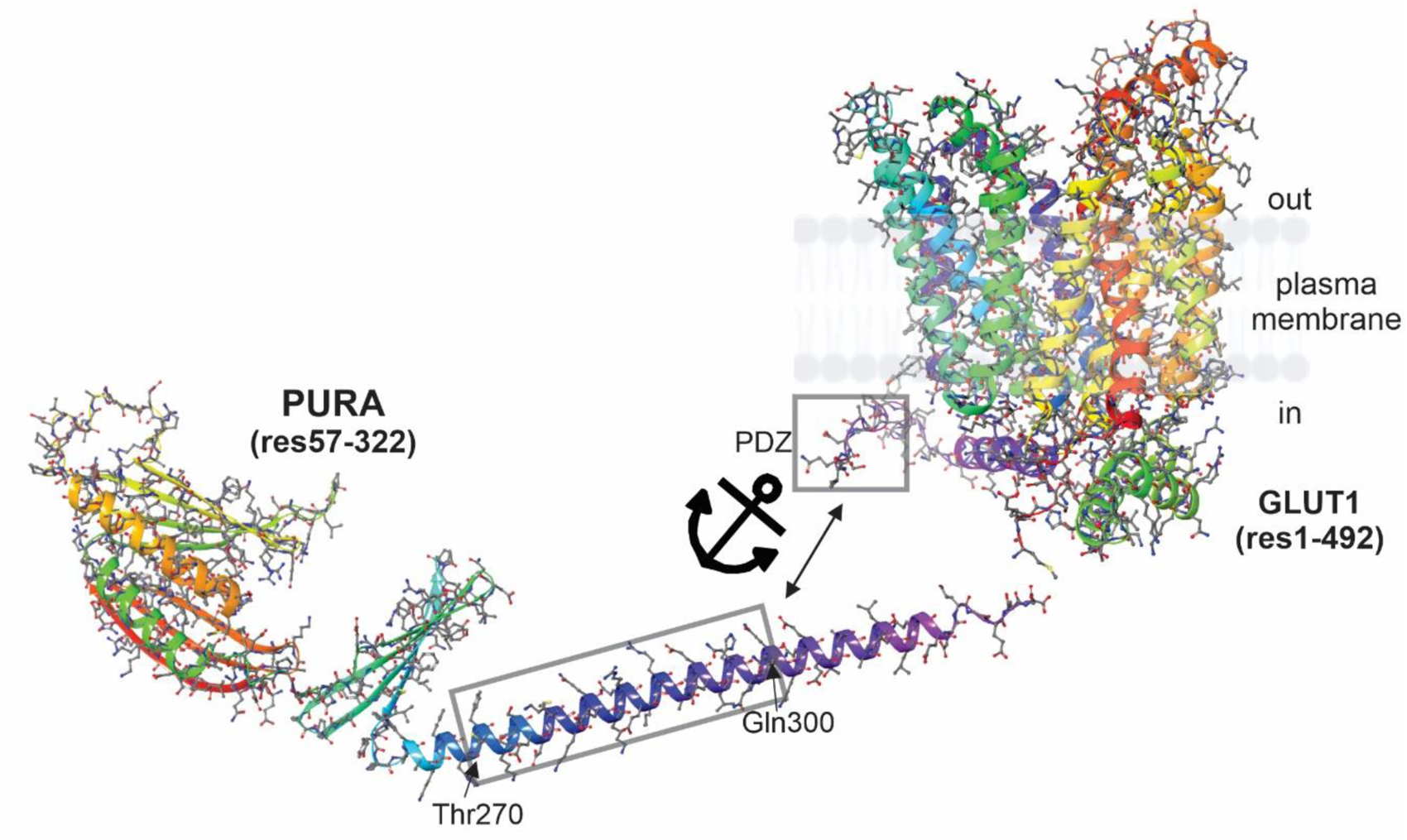
AlphaFold protein predictions. **Left**: wild-type PURA (res. 57-322). **Right**: GLUT1 including the PDZ binding motif (res 488-492, DSQV). Both proteins are rainbow-colored from N-terminal to C-terminal (from red to violet). Grey squares frame the regions from both proteins that were used as initial binding sites for docking purposes: PURA: Thr270-Gln300; GLUT1: PDZ motif: res. 488-492. Anchor + double-headed arrow is meant to point out the proposed docking region.

Protein-protein complexes were constructed via molecular docking with the ZDOCK server[55,56]and refined using atomistic molecular dynamics to account for full flexibility of the docking-generated PURA:GLUT1 complexes[57–59] using Gromacs-2023[41,60].

Four PURA versions (wild-type + 3 mutants) in complex with GLUT1 were studied, as detailed below (mutations are shown in red):

(i) wild-type PURA (res 57-322): ETQELASKRVDIQNKRFYLDVKQNAKGRFLKIAEVGAGGNKSRLTLSMSVAVEFRDYLGDFIEHYAQLGPSQPPDLAQAQDEPRRALKSEFLVRENRKYYMDLKENQRGRFLRIRQTVNRGPGLGSTQGQTIALPAQGLIEFRDALAKLIDDYGVEEEPAELPEGTSLTVDNKRFFFDVGSNKYGVFMRVSEVKPTYRNSITVPYKVWAKFGHTFCKYSEEMKKIQEKQREKRAACEQLHQQQQQQQEETAAATLLLQGEEEGEED
(ii) PURA-mut 1 (res 57-223): ETQELASKRVDIQNKRFYLDVKQNAKGRFLKIAEVGAGGNKSRLTLSMSVAVEFRDYLGDFIEHYAQLGPSQPPDLAQAQDEPRRALKSEFLVRENRKYYMDLKENQRGRFLRIRQTVNRGPGLGSTQGQTIALPAQGLSSSVTLWPSSSTTTEWRRSRPSCPRAPP
(iii) PURA-mut 2 (res 57-289): MADRDSGSEQGGAALGSGGSLGHPGSGSGSGGGGGGGGGGGGSGGGGGGAPGGLQHETQELASKRVDIQNKRFYLDVKQNAKGRFLKIAEVGAGGNKSRLTLSMSVAVEFRDYLGDFIEHYAQLGPSQPPDLAQAQDEPRRALKSEFLVRENRKYYMDLKENQRGRFLRIRQTVNRGPGLGSTQGQTIALPAQGLIEFRDALAKLIDDYGVEEEPAELPEGTSLTVDNKRFFFDVGSNKYGVFMRGEAHLSQLHHRALQGVGQVRTHLLQVLGGDEEDSREAEGEAGCL
(iv) PURA-mut 3 (res 57-322, Phe271 deleted): ETQELASKRVDIQNKRFYLDVKQNAKGRFLKIAEVGAGGNKSRLTLSMSVAVEFRDYLGDFIEHYAQLGPSQPPDLAQAQDEPRRALKSEFLVRENRKYYMDLKENQRGRFLRIRQTVNRGPGLGSTQGQTIALPAQGLIEFRDALAKLIDDYGVEEEPAELPEGTSLTVDNKRFFFDVGSNKYGVFMRVSEVKPTYRNSITVPYKVWA KFGHT_CKYSEEMKKIQEKQREKRAACEQLHQQQQQQQEETAAATLLLQGEEEGEED

Ranked complexes according to ZDOCK scoring function were visually inspected and a single protein-protein complex was selected for each one of the four PURA variants studied here. As a practical measure of protein-protein stability we estimated the total amount of potential donor-acceptor hydrogen bond pairs and the effective steady-state H-bonds formed at the protein-protein interface during the simulation[61,62], see Table 2.

**Table 2.** Protein-protein PURA variants: GLUT1 complex stability parameters: Predicted H-donors and acceptors plus established steady-state-H-bonds during simulation.

| PURA variant | Predicted H-donors | Predicted H-acceptors | steady-state H-bonds |
| --- | --- | --- | --- |
| wild-type | 1076 | 2087 | 33 |
| mut 1 | 939 | 1785 | 34 |
| mut 2 | 1022 | 1980 | 19 |
| mut 3 | 1075 | 2085 | 20 |

Table 2 suggests that PURA(wild-type) and PURA(mut 3) may establish the most amounts of interface H-Bonds, followed by PURA(mut 2) and PURA(mut 1), in agreement with the truncation lengths (at 289 and 223, respectively). However, the effective amount of hydrogen bonds that form within the protein-protein interface is significantly higher for PURA(wild-type) than for PURA(mut 3) even though PURA(mut 3) differs only in one deleted amino acid (Phe271). The protein-protein affinity is drastically reduced, giving the deleted Phe271 right at the beginning of the alpha-helix in the Repeat-III PURA region, a strategic location for the interaction with the GLUT1’s C-terminal PDZ binding domain. PURA(mut 2), which generates a mutated-truncated 289-amino acid version, reduces its affinity for GLUT1 in a third and interacts with GLUT1 amino acids between positions 445-451 (not the PDZ-binding motif).

PURA(mut.1) although truncated at 223, and even with the total absence of the Repeat-III region established as much as interface H-bonds as the wild-type protein, however, through PURA residues far away from the normally proposed PURIII-PDZ interaction (PURA-Glu57, Thr58, Asn80 and Lys82). Taking into account that the patient carrying mut 1 (Patient 1) clinically seems to have the least functional GLUT1, this abnormal and stable interaction could be adding a negative dominant effect for this mutation.

See figure 4 for final molecular dynamics snapshots and residues involved at each protein-protein interface.

**Figure 4.**
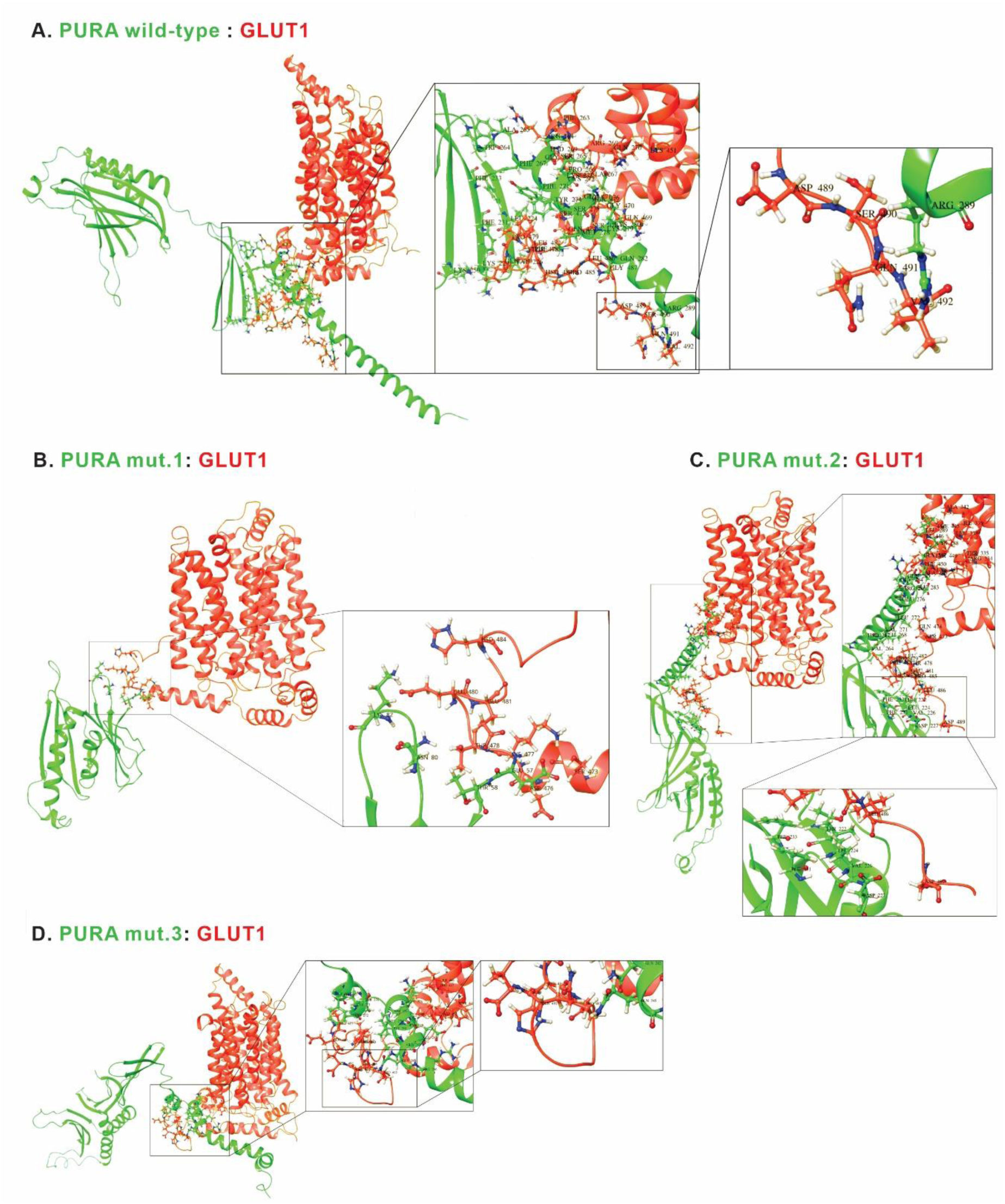
Molecular dynamics snapshots and residues involved at each protein-protein interface. PURA protein in green, GLUT1 in red. First zoom-in: PURA:GLUT1 interface, second zoom-in: PURA-GLUT1 PDZ interface. **4A**. PURA wild-type: GLUT1. PURIII repeat interacting with GLUT1’s PDZ binding domain, as expected. **4B**: PURA-mutant 1 (PUR repeat I) interacting with GLUT1(not involving PDZ binding motif). **4C**: PURA mutant 2 (abnormal PURIII) interacting with GLUT1’ PDZ binding domain. **4D**: PURA-mutant 3 (abnormal PURIII) interacting with GLUT1 (not involving PDZ binding motif). Corey–Pauling–Koltun coloring for atoms: white for hydrogen, blue for nitrogen, red for oxygen.

## DISCUSSION

Our study delves into the multifaceted roles of PURA, a protein with a well-established involvement in cellular processes across various cellular domains. PURA’s intricate functions have garnered considerable attention in the past years due to its association with neurological disorders and genetic syndromes.

The identification of hypoglycorrhachia as a consistent clinical feature in PURA syndrome, first reported by our team in 2018, raised questions regarding the molecular basis of this phenomenon. Here, we have extended our investigation to include a cohort of new patients exhibiting this hallmark symptom, shedding light on a potential regulatory role of PURA in glucose homeostasis through GLUT1.

Intriguingly, our functional experiments revealed a novel, fundamental role for PURA in cellular glucose uptake. The increase in glucose intake when PURA is upregulated (*no* different from GLUT1’s upregulation effect) and its diminishment upon PURA knockdown even in the presence of high GLUT1 levels suggests that GLUT1’s function strongly depends on PURA. These findings implicate PURA as a previously unrecognized factor in glucose transport, indicating its potential role in fine-tuning cellular glucose homeostasis and serves to explain the clinical observations in the 6 included PURA-patients described in this paper. While PURA’s well-established role as a transcriptional and translational modulator led us to consider an impact on GLUT1’s expression, our experiments provided evidence that this was not their way for interplay. Our work supports the idea of a PURA:GLUT1 complex formation to explain the their interdependent function.

Being GLUT1 a plasma membrane protein, with 12 transmembrane domains[52] its degradation and recycling depends on vesicular transportation. It has been described to enter the endocytic pathway and is recycled as a retromer-mediated cargo[49,63]. Knock-out of the well-established endocytic GTPase Rab21 disrupts GLUT1 recycling and decreases glucose uptake in HeLa cells[63]. Furthermore, GLUT1 has been shown to interact with Myosin 6 and the kinesin superfamily protein KIF-1B via its transporter binding protein GLUT1CBP (GLUT1 transporter binding protein) [64,65], probably having an important role in recycling of the glucose transporter and specific localization to the plasma membrane for its functioning. On the other hand, PURA is also known to closely associate with motor proteins[1]. The cargo transporting conventional kinesin KIF5[20] and myosin Myosin 5a[19,66] have been found to interact with PURA. Given these facts, one could speculate that PURA could play a role in GLUT1’s trafficking and specific localization for correct functioning. Location of glucose transporters on the plasma membrane are not random[67] and specially when one is talking about the blood-brain-barrier[38,68]. The implications of our results extend beyond the cellular context. We propose that the interplay between PURA and GLUT1 may have broader physiological consequences, potentially influencing glucose transport across the blood-brain barrier.

In conclusion, our study uncovers a previously unrecognized role for PURA in glucose uptake regulation and establishes a functional and physical interaction between PURA and GLUT1. These findings provide valuable insights into the molecular basis of hypoglycorrhachia in PURA-related neurodevelopmental disorder and suggest a potential regulatory role of PURA in glucose homeostasis.

Our results warrant further investigations into the specific cellular compartments of interaction and functional interplay between PURA and GLUT1. Additionally, it is imperative to explore the structural and functional implications of PURA mutations, particularly in the context of GLUT1 interactions.

## Acknowledgements

Firstly, we thank the patients and their families for contributing to research in the field. We also acknowledge: Dr. M. Rodriguez-Peña (Facultad de Odontología Universidad de Chile, reagent provision), Dr C. Tomes and Dr. M. Pavarotti (IHEM, Mendoza, Argentina, reagent provision), Dr. LS. Mayorga and Dr. M.Roqué (IHEM, Mendoza, Argentina – manuscript revision). Centro de Computación de Alto Desempeño de la Universidad Nacional de Córdoba (CCAD-UNC) for supercomputing time.

## Author contributions

LM conducted the research project. RC performed the wet lab experiments with the help of LM. DMas carried out the computational methods and analyses. CM, MA and DMar provided clinical data from patients. LM wrote the paper with the help of DMas for computational sections. All authors read and approved the final manuscript.

## Supplementary material

**Figure S1.**
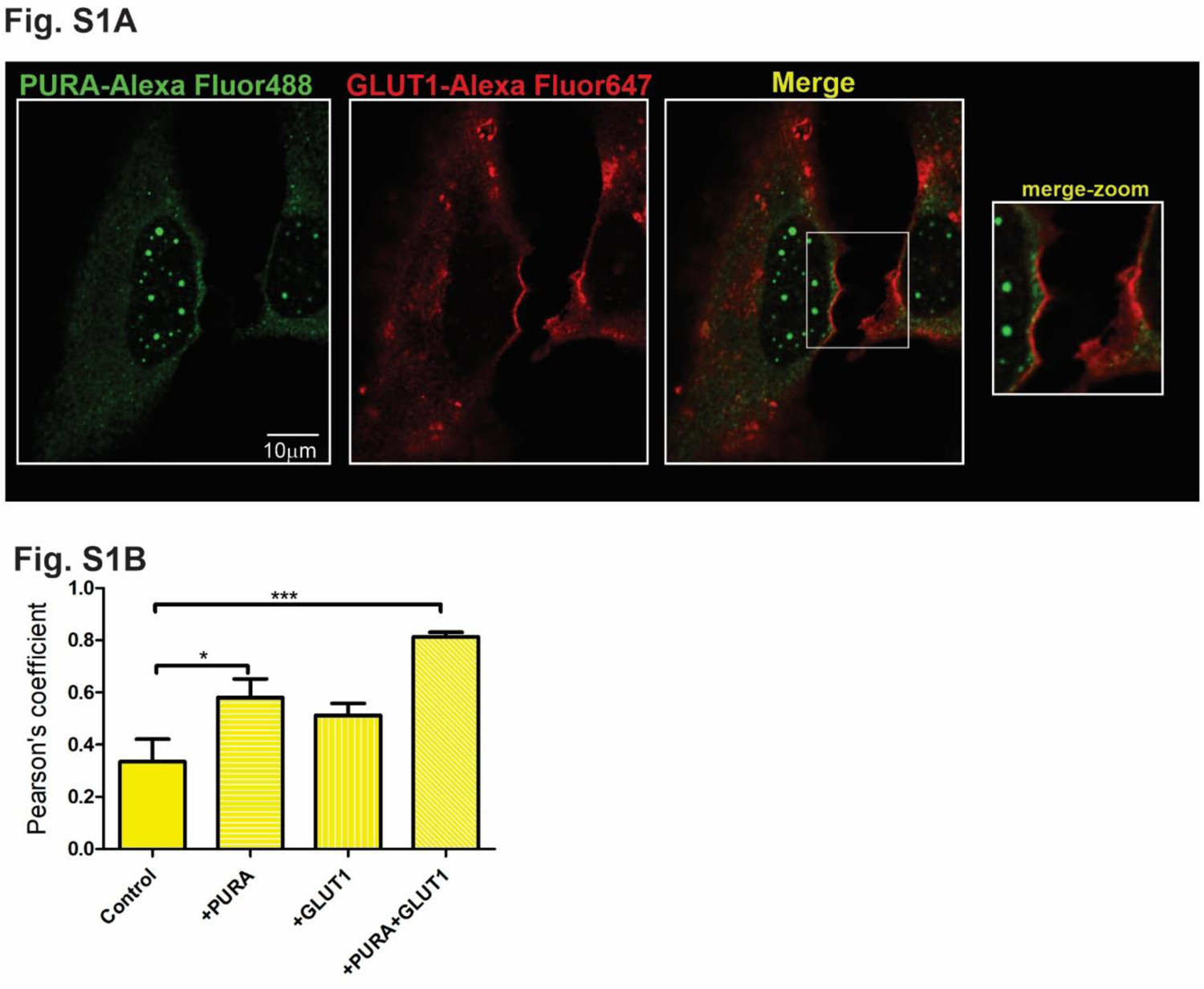
**S1A**: Confocal Immunofluorescence Microscopy. Confocal laser micrographs depict indirect immunofluorescence staining of PURA (labeled with AlexaFluor 488 in green) and GLUT1 (labeled with AlexaFluor 647 in red) within HeLa cells overexpressing PURA and GLUT1 expression. Colocalization of the two proteins in the merged image is represented as yellow, zoomed in a white square. **S1B**: Pearsońs coefficients values from 3 controls, 6 +PURA, 3 +GLUT1, 9 +PURA+GLUT1 photos (3+PURA +Rotenone, 6 +PURA+GLUT1). *** p<0.001, *p<0.05. One-way ANOVA+ Dunnett’s Multiple Comparison Test.

**Figure S2:**
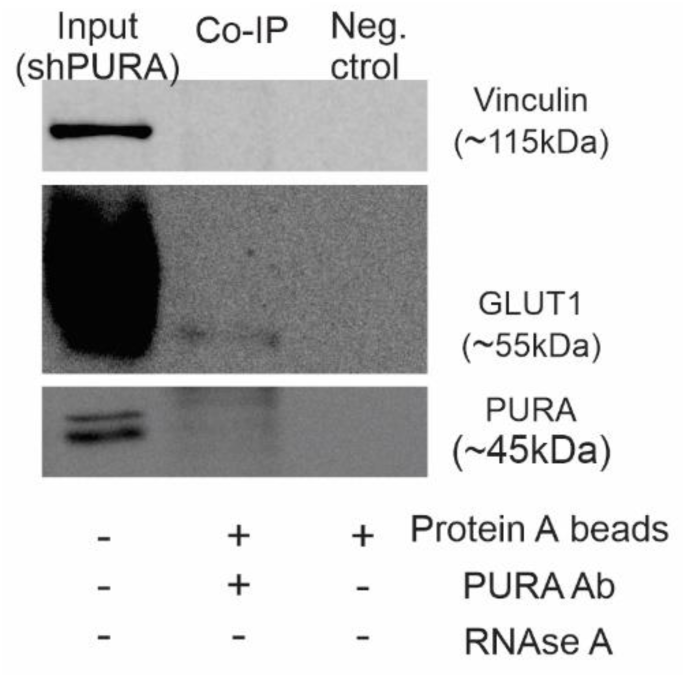
Co-immunoprecipitation of GLUT1 and PURA using Protein A agarose beads bond to anti-PURA antibody. shPURA input was used for the pull-down. There is presence of GLUT1 in the pull-down fraction of cells with down-regulated PURA. Anyhow, note that it appears when GLUT1 is overexposed, noting that a small fraction of GLUT1 is being able to bind to the scarce PURA protein.

